# Perceptual consequences of retinal stabilization with high-frequency non-stroboscopic displays

**DOI:** 10.1101/2024.05.02.592177

**Authors:** Y. Howard Li, Soma Mizobuchi, Jie Z. Wang, Michele Rucci

## Abstract

During natural fixation, ocular drifts continually modulate the input to the retina. Previous studies have shown that this motion enhances sensitivity to fine spatial detail, a conclusion supported by findings of reduced sensitivity to high—but not low—spatial frequencies when stimuli are immobilized on the retina for brief periods of time. Most prior retinal-stabilization studies have relied on fast-phosphor cathode ray tube (CRT) displays or adaptive optics scanning laser ophthalmoscopes (AOSLOs), both of which deliver temporally pulsed stimulation. This raises the question of whether stimulus flicker contributed to the previously observed perceptual impairments under retinal stabilization. Here, we replicate stabilization experiments using two types of fast displays that provide more continuous stimulation: liquid-crystal display (LCD) and organic light-emitting diode (OLED) monitors. We again find an impairment in sensitivity to high spatial frequencies under retinal stabilization. Analyses of the retinal input confirm high-quality stabilization within the temporal bandwidth of human vision. These results show that retinal-stabilization effects are robust across display technologies and are little affected by the specific dynamics of modern displays.

## Introduction

Humans are unaware that their eyes are always in motion, even when attempting to maintain steady gaze. During the brief intervals between rapid gaze shifts (saccades and microsaccades) a meandering, jittery motion—commonly known as ocular drift—continually shifts the image on the retina across many photoreceptors^1^ (Fig. 1A). The temporal modulations generated by this fixational motion provide computational advantages for encoding spatial information ^2–5^, and it has long been proposed that these signals contribute to visual processing^6–9^.

**Figure 1.**
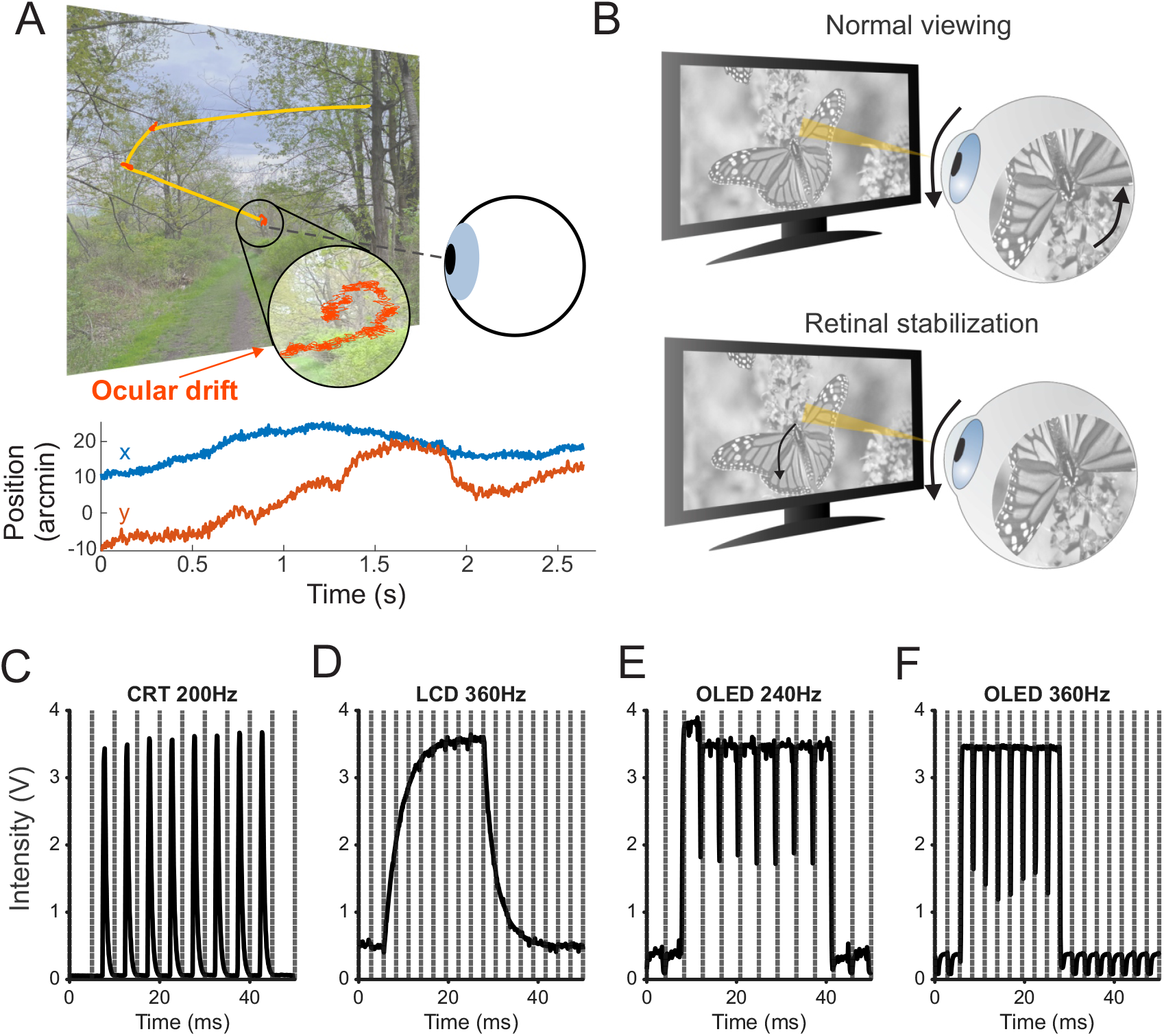
Eye movements, retinal stabilization, and display dynamics. **(A)** An eye movement trace (yellow) superimposed on the observed scene. Naturally occurring ocular drifts during inter-saccadic fixations are marked in red. The horizontal and vertical eye positions of the last fixation (zoomed-in inset) are shown in the bottom panel. **(B)** Retinal stabilization procedure. During normal viewing, eye movements cause incessant motion on the retina. Under retinal stabilization, the stimulus moves on the display to counteract the movement of the eye and maintain its projection at a fixed position on the retina. **(C–F)** Refresh dynamics of four types of monitors: a CRT at 200 Hz *(C)*, an LCD at 360 Hz *(D)*, and two OLEDs at 240 Hz *(E)* and 360 Hz *(F)*. Each monitor was driven to display a square gray patch at the center, and luminance was recorded with a photodiode, while the stimulus alternated between black and white every 8 frames (16-frame period). The vertical dashed lines mark the display refresh times.

Recent studies using retinal stabilization, a condition in which the stimulus moves together with the eye to remain stationary on the retina, have provided strong support to this idea: sensitivity to high-spatial-frequency stimuli is selectively impaired when retinal motion is eliminated for brief periods approximating the duration of natural inter-saccadic fixation, as expected if the visual system makes use of the luminance modulations delivered by fixational drifts^10–14^. In fact, spatial sensitivity closely follows the strength of the fixational modulations in experiments with precisely controlled retinal image motion^14^. Furthermore, ocular drifts are modulated in task-dependent manner consistently with the purposeful use of the resulting spatiotemporal signals^13,15^, and visual acuity varies systematically across individuals according to the characteristics of their ocular drifts.

While a growing body of evidence supports the view that fixational drift contributes to an active strategy for encoding visual information^4,12^, most retinal stabilization experiments have relied on systems that deliver *flickering* stimulation to the retina. For instance, cathode ray tube (CRT) displays render images sequentially, as an electron beam scans the screen line by line from top to bottom with each frame^16^. Similarly, adaptive optics scanning laser ophthalmoscopes (AOSLOs) generate retinal stimuli by turning a scanning laser on and off, typically at lower refresh rates than CRTs^11^. These presentation methods have raised the question of whether stroboscopic stimulation influenced the results of retinal stabilization experiments^17^.

To address this question, here we replicate previous retinal stabilization experiments using two types of modern displays: liquid crystal display (LCD) and organic light-emitting diode (OLED) monitors. Both technologies present more continuous stimulation, albeit through different mechanisms^18–20^. LCDs eliminate inter-frame blanks by employing a constant backlight and updating the voltages applied to the liquid crystals only when the image changes. OLEDs, in contrast, use self-emissive pixels that can be individually and rapidly updated with new image data, enabling near-instantaneous pixel refresh while maintaining constant illumination throughout each frame. For both display types, growing demand for high-performance graphics has led to the development of monitors with high refresh rates, which can, in principle, be used to precisely control retinal stimulation during eye movements.

## Results

In retinal stabilization experiments, the position of the stimulus is updated at each refresh of the display to compensate for the observer’s eye movements, thereby keeping the stimulus stationary on the retina (Fig. 1B). To minimize errors caused by fast eye movements, this procedure is typically restricted to periods of relatively slow intersaccadic fixation, while trials containing saccades or microsaccades are excluded from analysis. In this study, we first evaluate the quality of retinal stabilization resulting from distinct display technologies. We then present psychophysical measurements obtained using different monitors to examine possible influences of display dynamics.

### Quality of retinal stabilization

Fig. 1C-F illustrates the temporal luminance dynamics of four displays—a CRT, an LCD, and two OLED monitors—when displaying a uniform 25 mm × 25 mm square patch. The luminance signals were recorded using a photodiode (Vishay Intertechnology Inc.) while the patch alternated between black and white in a square-wave modulation with a 16-frame period. Each display was tested at its maximum refresh rate: 200 Hz for the CRT, 360 Hz for the LCD, and 240 Hz and 360 Hz for the two OLEDs, respectively.

As expected, the CRT monitor exhibits the stereotypical stroboscopic pattern, with brief luminance spikes occurring at every passage of the beam interleaved by long blank periods as the phosphors decay (Fig. 1C). In contrast, the LCD monitor gives a smoother temporal profile, with gradual increases and decreases in luminance and no interruptions in visual stimulation (Fig. 1D). The two OLED monitors (Fig. 1E–F) are characterized by fast response times and steady luminance levels, except for brief transient dips in intensity immediately before each refresh of the image due to so-called inter-frame blanking, a common feature of OLED monitors. Unlike the CRT, however, these interruptions have extremely short durations, lasting less than 0.5 ms.

To estimate the quality of retinal stabilization afforded by each monitor, we modeled the changes in luminance that a retinal photoreceptor would experience in a typical experiment (Fig. 2A). Based on the photodiode measurements, we derived temporal filters that captured the persistence and temporal response of each monitor. Specifically, we used a gamma kernel to model the luminance spikes of the CRT monitor and first-order low-pass filters with separate time constants for luminance increments and decrements for the LCD and OLED displays, and we tuned the respective parameters to closely fit the recorded luminance data (Fig. 2B). We modeled fixational eye drift as a Brownian motion process with diffusion constant *D* = 20 arcmin^2^/s, a model that well replicates the characteristics of inter-saccadic eye movements^21–23^. As in previous retinal stabilization experiments with stimuli at high spatial frequencies^10,12^ and the experiments described later in this article, we assumed the stimulus to be a 10 cycles/deg grating.

**Figure 2.**
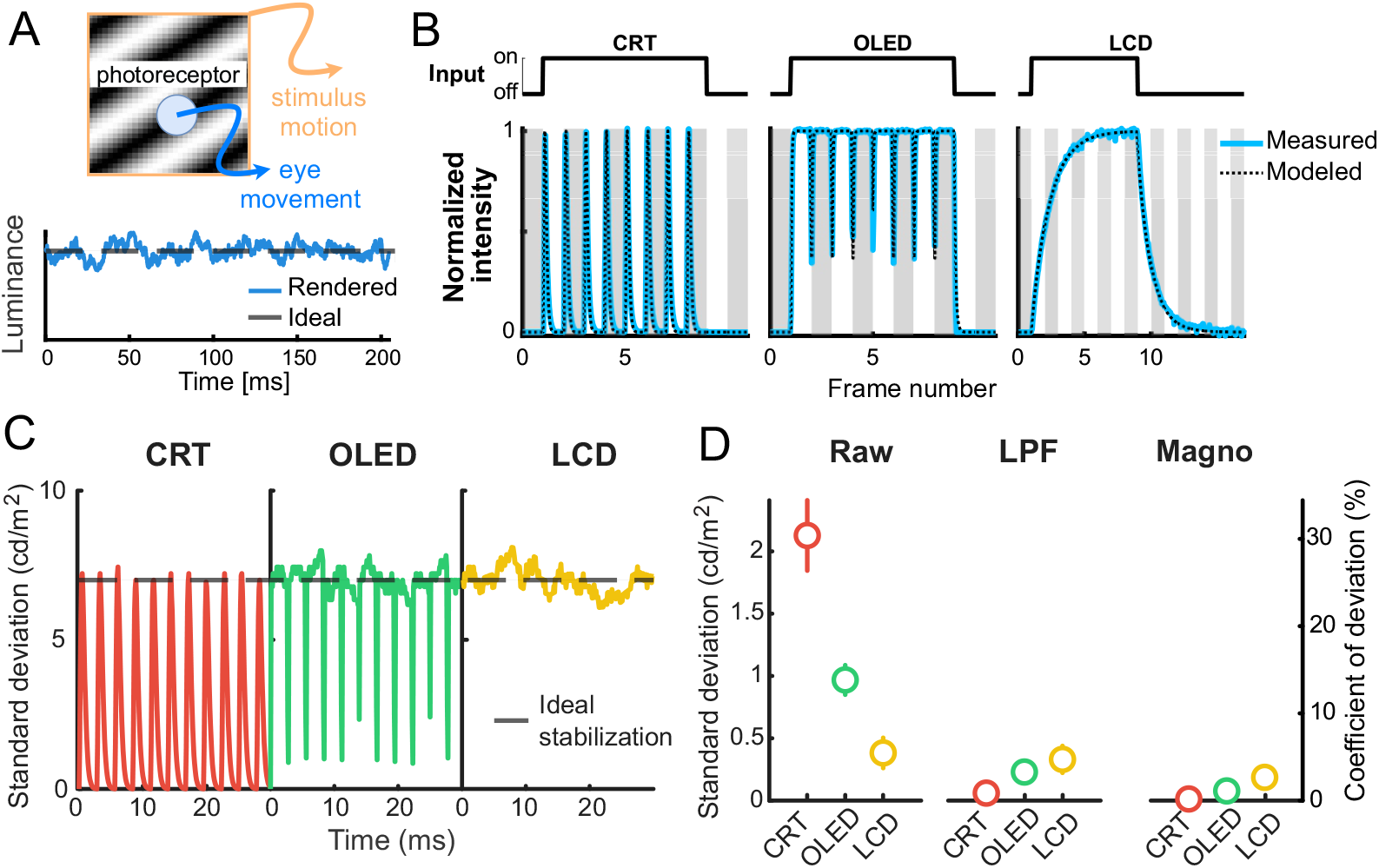
Impact of display refresh dynamics on retinal stabilization. **(A)** Ideally, compensating for the observer’s eye movements (blue trace) by continually updating the stimulus on the monitor (orange trace) should produce a constant visual input at the level of individual photoreceptors (bottom panel; dashed line). In practice, however, even with perfect oculomotor measurements, the temporal dynamics of the display introduce fluctuations in the input signal (bottom panel; blue trace). **(B)** Modeling monitor dynamics. The temporal characteristics of three monitors were modeled based on the luminance data measured while driving each monitor with a 16-frames square wave. Monitors were driven a_**C**_t _**R**_th_**T**_eir maximum frequencey: 200 Hz for the CRT (left), 360 Hz for the OLED (middle) and for the LCD (right). For each monitor, the simulated intensity values (dashed black lines) are compared to the luminance measurements (solid blue lines). **(C)** Simulated retinal input during gaze-stabilized fixations. The gray dashed line represents the case of sideal stabilization with unchanging retinal input. Colored lines show the modeled luminance signals experienced under retinal stabilization with the CRT (left, red), the OLED (middle, green), and the LCD (right, yellow). **(D)** Estimated variability in the retinally stabilized input stimulation provided by the three monitors. Data points represent the standard deviation (left ordinate) and corresponding coefficient of variation (right ordinate) for the raw input signals (left panel), after low-pass filtering at 80 Hz (center panel), and after filtering by the temporal sensitivity of magnocellular ganglion cells (right panel). Errorbars represent ± one standard deviation across 20 simulated photore-ceptors and 50 fixations.

Under perfect stabilization, the input signal experienced by a photoreceptor would remain constant over time (dashed lines in Fig. 2A and Fig. 2C), as the stimulus would move to counteract the consequences of eye movements. In practice, however, luminance varies because of the characteristics of each display, which result in temporal fluctuations in the input signal (colored lines in Fig. 2C). These modulations differ across display types: they contain gradual changes when the stimulus is displayed on the LCD (yellow trace in Fig. 2C), brief luminance pulses for the CRT (red trace), and extremely brief interruptions in visual stimulation for the OLEDs (green trace).

We evaluated the precision of retinal stabilization by calculating how much the luminance input to any given retinal location deviates from the ideal case of constant stimulation (Fig. 2D). As expected, when looking at the raw input data, the luminance pulses introduced by stroboscopic displays, such as CRTs, cause large deviations from the intended values. The other monitors did not exhibit the large deviations present in the CRT because of their more sustained responses. This was particularly evident for the LCD, which delivers almost constant stimulation once it reaches its steady state. To quantify the quality of retinal stabilization, the left panel of Fig. 2D shows the standard deviation (left ordinate) and the corresponding coefficient of variation (the percentage deviation from the mean, right ordinate) experienced by the modeled photoreceptor in a typical trial of a retinal stabilization experiment for the three types of monitors. In terms of the full (unfiltered) luminance signals, the LCD monitor yields the smallest variability, with about a 5% change in luminance during the course of an experimental trial.

It is important to emphasize that the data reported in the left panel of Fig. 2D provide an overestimation of the effective error in retinal stabilization. These data consider all fluctuations in the input signal at all temporal frequencies, irrespective of whether they are within or outside of the temporal sensitivity bandwidth of the visual system. For example, much of the deviation in the CRT occurs at frequencies well beyond the visual system’s sensitivity. To elucidate this, consider an ideal CRT display that instantaneously updates phosphors with zero persistence. In this case, the input stimulus would be a train of flashes separated by the refresh interval (in our case 5 ms, since the monitor was updated at 200 Hz), which in the frequency domain would correspond to a stack of harmonics at integer multiples of the display frequency. Only the DC (0 Hz) component of this stimulus would be detected by the visual system (corresponding to our percept of an unflickering monitor), because all the other harmonics are well beyond what can be resolved by the visual system. Considering very high frequency harmonics leads to an inflated estimation of the perceptual impact of the stabilization error.

To estimate the perceptually relevant consequences of imperfect stabilization, we examined the stabilization error after filtering the input signals by the temporal sensitivity of human vision. The data in the center and right panels of Fig. 2D quantify the fluctuations in luminance experienced by the simulated photoreceptor after applying two perceptually relevant filters: a low-pass filter with an 80 Hz cutoff frequency, a conservative upper limit for flicker fusion^24,25^ (Fig. 2D, center panel); and a filter designed to replicate the temporal of macaque magnocellular ganglion cells—the neurons in the retina sensitive to higher temporal frequencies—-, as measured in neurophysiological recordings^26^ (Fig. 2D, right). Both analyses show that all monitors provide high-quality of retinal stabilization within the relevant temporal bandwidth. Notably, the ranking across displays changes after filtering, as rapid interruptions—such as the stroboscopic rendering of the CRT and the inter-frame blanking of the OLED—are largely beyond the range relevant for perception and for driving neural responses. Accordingly, the CRT monitor becomes the one with the highest quality of retinal stabilization, with input variability smaller than 1%.

### Perceptual consequences of retinal stabilization

Having established that all three monitors yield high-quality of retinal stabilization, we examined whether elimination of retinal motion with OLED and LCD displays for short periods of time impairs discrimination of fine patterns, as previously reported with CRT displays.

The experimental procedure was similar to our previous experiments with retinal stabilization^10,12,14^. In a forced-choice paradigm, subjects were asked to report whether a high-frequency grating was oriented by 45° to the left or to the right (Fig. 3A). Each trial started with the observer fixating on a marker at the center of a uniform gray field. To minimize luminance transients caused by stimulus presentation, the stimulus, with orientation that varied randomly across trials, then appeared gradually with linearly increasing contrast over an initial period (800 ms), which was followed by 500 ms exposure at the plateau contrast. A large-field maximum contrast noise mask ended the trial, prompting subjects to report the perceived orientation of the grating by pressing one of two buttons on a joypad (Fig. 3B).

**Figure 3.**
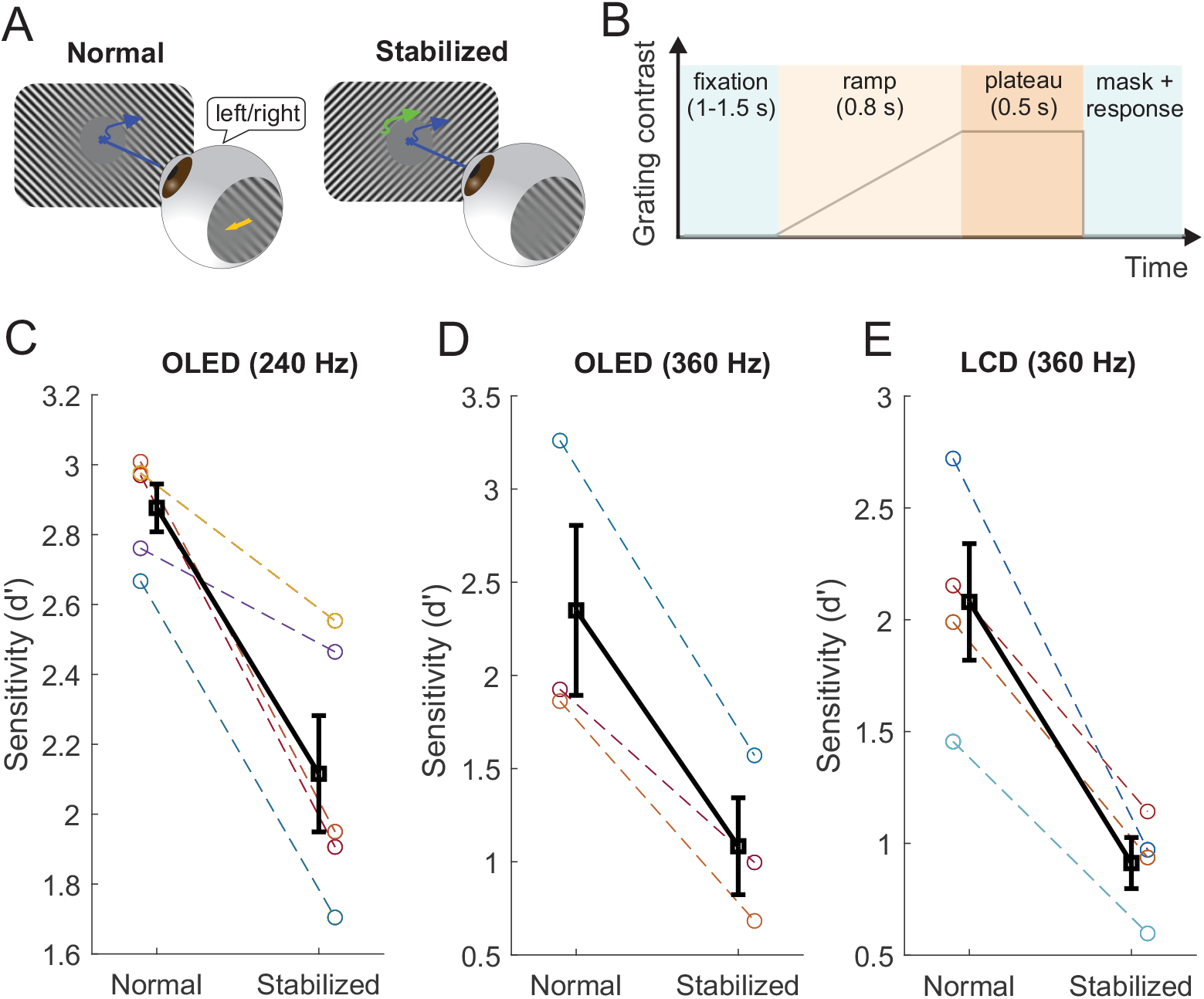
Perceptual consequences of retinal stabilization. **(A)** A high-frequency orientation discrimination task under normal viewing (left) or retinal stabilization (right). **(B)** The temporal sequence of a trial. Stimulus contrast ramped up for 0.8 s followed by a 0.5 s plateau period. A full-screen noise mask ended the trial. **(C–E)** Measured sensitivity (expressed as *d*^′^) when stimuli were rendered on three different monitors: **(C)** OLED monitor at 240 Hz. Stabilizing the stimulus on the retina impairs sensitivity for all subjects (*N*=5; *p*=0.01, paired *t*-test). **(D)** OLED monitor at 360 Hz (*N*=3; *p*=0.03, paired *t*-test). **(E)** LCD monitor at 360 Hz (*N*=4; *p*=0.01, paired *t*-test). In all panels, thin lines indicate individual subjects and the black line shows the average ± SEM across subjects. Across all display types, retinal stabilization reduced sensitivity for high-frequency orientation discrimination.

Trials randomly alternated between two conditions: normal viewing and retinal stabilization. In both conditions, eye movements were continually recorded by means of a high-resolution dual Purkinje Image eye-tracker^27^, a system that provides arcminute resolution with ∼1 ms delay. Following the approach used in our previous studies, stabilization was achieved by means of a custom system for gaze-contingent display control^28^. This approach has been shown to achieve high-quality retinal stabilization with errors of the order of one arcminute^10^. As customary in retinal stabilization experiments, to isolate the influences of the luminance modulations resulting from ocular drift, all trials containing blinks, saccades, or microsaccades during stimulus presentation were excluded from the analysis. In separate experimental sessions, distinct groups of subjects were exposed to stimuli rendered on the three monitors, an LCD and an OLED driven at 360 Hz (both modeled in Fig. 2), and an OLED driven at 240 Hz (see Fig. 1*C*–*F*; the).

As shown in Fig. 3C–E, retinal stabilization greatly impaired fine pattern discrimination, irrespective of the type of display used to render the stimuli. Similar decrements in performance were measured with the 240 Hz OLED (N = 5; p = 0.01, paired t-test; Fig. 3C), the 360 Hz OLED (*N* = 3; p = 0.03; Fig. 3D), and the LCD monitor (N = 4; p = 0.01; Fig. 3E). On average across subjects, sensitivity decreased by 43%, an impairment consistent with previous observations obtained using CRT displays at lower refresh rates. The reduction was consistent across observers, with every participant showing lower *d*^′^ under stabilization. These data show that the perceptual impairments previously reported under brief periods of retinal stabilization are robust with respect to display technology and refresh rate. Perception of high spatial frequency patterns is impaired in the absence of the visual input changes normally resulting from eye movements.

## Discussion

During natural fixation, the eye moves incessantly, continually translating the image on the retina and transforming a static scene into temporal luminance modulations at the level of individual photoreceptors. A growing body of work indicates that these luminance modulations contribute to spatial vision, particularly for fine detail. Notably, sensitivity to and discrimination of high—but not low—spatial frequencies is impaired when retinal motion is eliminated for brief intervals comparable to inter-saccadic fixation^10,11,13^. The present study extends these findings by testing a methodological concern: whether perceptual impairments reported under retinal stabilization could be attributed, at least in part, to the stroboscopic nature of the displays traditionally used to implement gaze-contingent stimulation (*e*.*g*., CRTs and adaptive optics scanning laser systems). Using high-refresh LCD and OLED monitors that provide more continuous stimulation, we again observed a robust reduction in sensitivity for high-spatial-frequency orientation discrimination when retinal image motion was removed. These results cannot be attributed to CRT flicker and strengthen the conclusion that stabilization-induced deficits reflect the functional role of ocular-drift–driven retinal modulations.

Our analyses include consideration of both photometric measurements (Fig. 1) and simulations of photoreceptor exposure under retinal stabilization (Fig. 2). Although CRT, LCD, and OLED monitors provide comparable image quality from a perceptual perspective, their physical luminance time courses differ markedly. In particular, in CRTs, stabilization errors computed from the unfiltered luminance signals are dominated by high-frequency components generated by stroboscopic rendering, which are, however, invisible to the visual system. Thus, direct examination of the raw signals overweights temporal fluctuations that are not perceptually relevant. Within the temporal bandwidth of visual perception and/or early neural responses, the dark period caused by CRT contribute minimal error. As a consequence, the effective quality of retinal stabilization of CRTs is even slightly higher than that of LCD displays, which do not employ stroboscopic rendering but tend to have slightly slower dynamics. In practice, all three monitor types yield high-quality retinal stabilization. High refresh rates further limit residual imperfections in gaze-contingent rendering by reducing the retinal displacement that can accumulate between stimulus updates. For example, with a typical drift speed of ∼30 arcmin/s and a 360 Hz update rate, the eye moves only ∼5 arcsec between refreshes—about 1.4% of the period of a 10 cycles/deg grating, corresponding to an average luminance change of ∼5.5% of the modulation amplitude. Any slower drift would reduce this bound proportionally. Thus, under the refresh rates and response dynamics tested here, the effective departures from ideal stabilization are minimal in the perceptually relevant domain. Displays with slower response times or lower refresh rates would be expected to yield larger residual modulations and should be evaluated using the same approach.

The reduction in performance measured when high spatial frequency stimuli are examined during brief periods of retinal stabilization suggests that eliminating the temporal luminance modulations normally resulting from ocular drifts impairs fine pattern vision. Neurons in the visual system are known to be highly sensitive to temporal changes^29–31^. Although this preference for non-zero temporal frequencies is typically examined with respect to the motion of objects in the scene, neurons are always exposed to temporal changes on the retina, as the self-motion of the observer generates dynamic sensory signals even for stationary objects—*i*.*e*., it redistributes the power of a static stimulus across nonzero temporal frequencies—offering valuable information about its spatial structure. The idea that the visual system uses temporal modulations to encode spatial patterns dates back more than a century^6,7^. Early proposals focused specifically on the limits of visual acuity, but using temporal changes from ocular drifts for encoding fine spatial patterns in general makes sense from a computational perspective, as the power that ocular drift makes available in the form of temporal changes (*i*.*e*., the power at non-zero temporal frequencies) increases with spatial frequency within a range that, in diffusion models, varies inversely with the drift diffusion constant^3^. During viewing of natural scenes, this reformatting counterbalances the power spectrum of natural scenes, yielding an effective visual input that equalizes (“whitens”) temporal power over a broad spatial frequency range, an operation that compresses information and has long been proposed as one of the functions of early visual processing^32,33^.

In keeping with this space-time encoding idea, it has been shown not only that eliminating drift-induced motion selectively impairs sensitivity to high spatial frequencies^10,11^ and that individual drift characteristics predict acuity limits^34^, but also that sensitivity closely follows the strength of the resulting modulations when retinal image motion is experimentally controlled^14^. Furthermore, humans adjust drift in a task-dependent manner in ways that enhance these modulations^13,15^. Importantly, the use of oculomotor-induced temporal transients is not limited to drift. Perceptual enhancement consistent with the spatial information conveyed by movement-generated transients has been reported for smooth pursuits^35^, microsaccades^36^, saccades^37^, and even blinks^38^. Although these behaviors differ markedly in their kinematics, they redistribute stimulus power across temporal frequencies following qualitatively similar profiles, with a progressively narrower whitening band that approaches zero for eye blinks. These results suggest a parsimonious active-encoding strategy in which common neural mechanisms operate on oculomotor-driven luminance transients generated by multiple types of oculomotor activity to support spatial vision. In this view, visual representations depend not only on the spatial pattern on the retina but also on how eye movements structure the incoming flow of luminance signals over time^3,14^.

In sum, our study shows that high-refresh CRT, LCD, and OLED displays yield comparable retinal stabilization quality and convergent perceptual consequences. More broadly, we provide a general approach for testing and validating gaze-contingent stimulation on modern display hardware. These findings establish high-refresh LCD/OLED monitors as reliable platforms for psychophysical and neurophysiological studies that probe the consequences of disrupting visuomotor loops via retinal stabilization.

## Methods

### Participants

Ten participants took part in this study (five males and five females; age range: 22–35). Five subjects participated in the OLED 240 Hz experiment (Fig. 3C), three in the OLED 360 Hz experiment (Fig. 3D), and four in the LCD experiment (Fig. 3E), with two participants completing both the OLED 360 Hz and LCD experiments. All participants were emmetropic and were compensated for their participation. With the exception of one author, all participants were naïve about the purpose of the experiments. Informed consent was obtained from all participants. The study was conducted in accordance with procedures approved by the University of Rochester Institutional Review Board.

### Stimuli and Apparatus

In three separate experiments, stimuli were displayed on an LCD at 360 Hz (ASUS PG259QN), an OLED at 240 Hz (LG UltraGear), and an OLED at 360 Hz (Alienware AW2725DF) in a dimly illuminated room. All three monitors were calibrated to linearize the relationship between input RGB intensity values and output luminance. Stimuli consisted of full-field sinusoidal gratings (24° × 15° for the LCD; 41° × 24° for the OLEDs) at high spatial frequency (16 cycles/deg on the LCD; 10 cycles/deg on the OLEDs) tilted by ±45° relative to the horizontal meridian and displayed over a uniform background.

To minimize luminance transients on the retina that were not caused by eye movements, the stimulus contrast ramped up linearly before reaching an individually selected value (Fig. 3B). Stimuli were either displayed at a fixed location of the monitor (normal viewing condition) or moved on the display to counteract the motion of the eye (retinal stabilization). To minimize the possibility that the observer would attempt to pursue the stabilized stimulus, thereby altering eye movements between the two conditions, foveal vision was eliminated by an artificial scotoma: a circular opaque gray region (1 ° in LCD; 7.5 ° in OLED) with a surrounding 0.5° annulus of tapered transparency, which remained centered on the line of sight as the eyes moved normally during fixation.

As in previous studies^13^, stimuli were viewed monocularly with the right eye while the left eye was patched. The participant’s head was immobilized by a dental imprint bite-bar and a headrest to maintain a fixed distance from the monitor (123 cm from the LCD and 78 cm from the OLEDs). Movements of the viewing eye were measured by a custom digital Dual Purkinje Image (DPI) eye-tracker^39^, a system with demonstrated sub-arcminute resolution^27^. Stimuli were rendered using EyeRIS, a system for gaze-contingent display control that enables real-time synchronization between oculomotor measurements and image updating on the monitor^28^ (average delay: 1.5 frames).

### Psychophysical Procedures and Data Analysis

Data were collected in several experimental sessions, each lasting approximately 1 hour. Each session was composed of several 10-15 minute blocks of trials, separated by breaks to allow participants to rest. Every session began with a two-step calibration procedure to ensure optimal eye-tracking and gaze-contingent control, as described previously^40^. In this procedure, the spatial registration between eye movements and the stimulus on the monitor was refined in a gaze-contingent phase in which the observer manually corrected possible offsets in the estimated position of gaze, shown in real time as a marker on the display.

The calibration was followed by blocks of experimental trials, in which participants reported whether the grating was tilted to the left or the right. Each trial started with the observer fixating on a center marker (a 10^′^ dot). After stable fixation was detected, the trial sequence described in Fig. 3B was initiated, with the contrast of the stimulus first ramping up for 800 ms and then remaining constant for 500 ms at a contrast value individually selected to yield ∼85% correct responses in the normal viewing condition. A full-field Brownian noise mask ended the trial, and prompted the observer to report the perceived orientation of the grating by pressing a corresponding button on a joypad. Both the orientation of the stimulus and the viewing condition—normal viewing (stimulus stationary on the monitor) or retinal stabilization (stimulus stabilized on the retina)—were randomly interleaved across trials.

The sampled oculomotor traces were segmented into periods of saccades and drifts based on instantaneous speed threshold of 2 °/s, as in our previous studies. Performance (reported as *d*^′^ in Fig. 3*C*–*E*) was evaluated on the trials in which the eye moved exclusively due to ocular drift during stimulus presentation. All trials containing blinks, saccades, or microsaccades were discarded, as these movements introduce additional luminance transients that are outside the scope of this study.

### Retinal Stabilization Assessment

Retinal stabilization quality was assessed by quantifying the residual retinal-image motion and the resulting luminance fluctuations on the retina for each display technology. Using photometric measurements of each monitor’s temporal response (Fig. 1), we modeled the luminance signal experienced by a simulated photoreceptor during fixation and under retinal stabilization (Fig. 2).

Monitor dynamics were measured by displaying flickering gray patches at the center of each monitor and acquiring the resulting luminance signals using a photodiode. For all monitors, contrast and brightness were both set to 50%, with all other settings set to factory defaults. The luminance of the patch was modulated as a square-wave with a duty cycle of 50% and a period of sixteen frames, an interval long enough to enable the LCD—the monitor with the slowest dynamics—to reach steady state within each half period. The modulation was set to 50% Michelson contrast and rendered using EyeRIS^28^. The photodiode (Vishay Intertechnology Inc.) was placed 1 cm from the display surface over the patch with a custom enclosure to limit light leakage outside the 25.4×25.4 mm region. The photocurrent was first converted into voltage by a high-bandwidth transimpedance amplifier, with gain values tuned for each monitor to maintain the output signal within [0–5] V. Analog voltages were then sampled at 10 kHz and quantized to 16-bit via a PCIe ADC board (General Standards Corporation), yielding traces like those reported in Fig. 1C-D.

Based on the recorded luminance traces, we modeled each monitor’s dynamics to derive temporal filters that simulated the output luminance 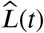 given a time-series of requested luminance values:

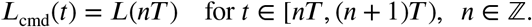

where *L*(*nT*) is the desired luminance value at each frame (*T* is the frame interval, *i*.*e*., the inverse of the refresh rate). For all monitors, luminance traces were modeled at 10 kHz (Δ*t* = 100 *μ*s) to match the sampling rate at which data were acquired.

The output of the CRT was estimated as

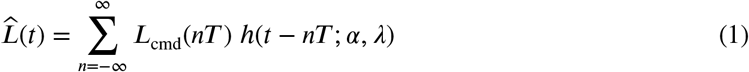

where *T* was 5 ms and *h*(*t*; *α, λ*) is a gamma-shaped kernel fitted to model the CRT pulses:

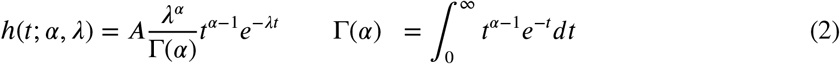

The parameters *α* = 3.196 and *λ* = 4.810^3^ s^−1^ were estimated to fit the photodiode measurements, and *A* was set to ensure that the kernel reached unit value at its peak 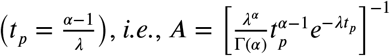.

The dynamics of the LCD and OLED monitors were modeled with a first-order difference equation:

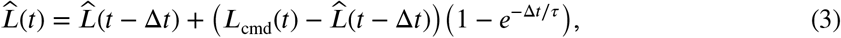

where two different time constants, τ_rise_ and τ_fall_, were used for increments or decrements in intensity, respectively. Time constants were obtained by fitting the luminance traces recorded from each monitor while ignoring inter-frame blanking in the OLED (the extremely brief luminance drops right before each new frame). Estimated parameters were: τ_rise_ = 3.7 ms and τ_fall_ = 2.8 ms for the LCD, and τ_rise_ = 0.8 ms and τ_fall_ = 0.2 ms for the OLED.

To model the OLED intra-frame blanking, the output luminance 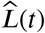 from Eq. 3 was multiplied by a periodic gating function *p*(*t*):

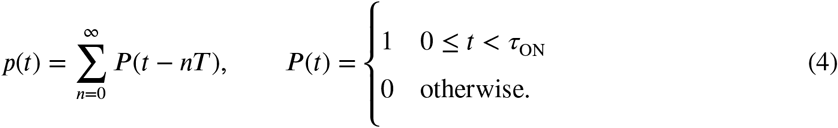

where τ_ON_ is the emissive period, fit to the measured luminance dynamics (τ_ON_ = 2.6 ms).

Retinal stabilization errors were estimated by modeling the luminance fluctuations that would be experienced by an individual photoreceptor as a consequence of the measured display dynamics and gaze-contingent updating procedures. Specifically, ocular drift was modeled as a Brownian motion process with diffusion constant *D* = 20 arcmin^2^/s, a realistic value for high-acuity stabilization experiments. Oculomotor traces were generated at 10 kHz to match the time step of the temporal filters in Eqs. 1–4, and were then downsampled to 1 kHz and quantized to pixel coordinates (1 pixel = 6^′^) to emulate the processing pipeline of a typical retinal stabilization experiment. The stimulus was a 10 cycles/deg grating.

For each monitor, we modeled the luminance experienced by an array of 20 photoreceptors equispaced over one period of the grating, as the eye drifted for 500 ms. Each photoreceptor was assumed to only cover 10 arcseconds to provide a conservative upper estimate of the stabilization error. We quantified the degree to which retinal input deviates from ideal stabilization by computing the standard deviation and the corresponding coefficient of variation of the simulated raw luminance signals as well as of the signals filtered by two perceptually motivated temporal filters: a fourth-order Butterworth IIR low-pass filter with an 80 Hz cutoff frequency, a conservative upper limit for human achromatic flicker fusion^24,25^; and a filter simulating the responses of macaque magnocellular retinal ganglion cells^41^:

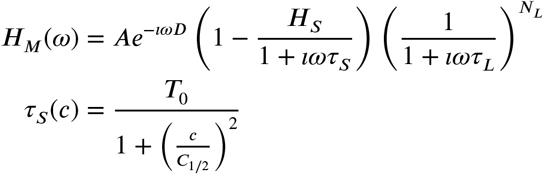

This model was originally developed for cat X cells and later shown to well predict also the responses of macaque P and M cells with proper tuning of the parameters^42,43^. Parameters were taken from the neurophysiological literature to model the responses of cells located at approximately 5° of eccentricity^43^: *N*_*L*_ = 30.3, *A* = 499.77 impulses ⋅ s^−1^, *D* = 2.0 ms, *H*_*S*_ = 1.00, τ_*L*_ = 1.44 ms, *T*_0_ = 37.34 ms, *C*_1/2_ = 0.051, *c* = 0.50. The data in Fig. 2*D* reflect averages across 1,000 simulated luminance traces (20 photoreceptors × 50 fixations) per monitor.

## Acknowledgments

This work was supported by grants R01 EY18363 and P30 EY001319 from the National Institutes of Health. We thank the members of the Active Perception Laboratory at the University of Rochester for helpful discussions.

## Author contributions

YHL and MR conceived the study. YHL, JZW, and SM implemented the psychophysical experiments, collected and analyzed the behavioral data. SM implemented the monitor measurements, collected and analyzed the measurement data. All authors contributed to the interpretation of the results and the writing of the article. MR supervised the project.

